# Systemic antisense therapeutics inhibiting *DUX4* expression improves muscle function in an FSHD mouse model

**DOI:** 10.1101/2021.01.14.426659

**Authors:** Ngoc Lu-Nguyen, Alberto Malerba, George Dickson, Linda Popplewell

## Abstract

Aberrant expression of the double homeobox 4 (*DUX4*) gene in skeletal muscle causes muscle deterioration and weakness in Facioscapulohumeral Muscular Dystrophy (FSHD). Since the presence of a permissive pLAM1 polyadenylation signal is essential for stabilization of *DUX4* mRNA and translation of DUX4 protein, disrupting the function of this structure can prevent expression of DUX4. We and others have shown promising results using antisense approaches to reduce *DUX4* expression *in vitro* and *in vivo* following local intramuscular administration. Our group has developed further the antisense chemistries, and demonstrate here enhanced *in vitro* antisense efficacy. The optimal chemistry was conjugated to a cell-penetrating moiety, and for the first time in FSHD research has been systemically administered into a double-transgenic mouse model of FSHD. After four weekly treatments, mRNA quantities of *DUX4* and target genes were reduced by 50% that led to a 5% increase in muscle mass, a 52% improvement in *in situ* muscle strength, and reduction of muscle fibrosis by 17%. Systemic DUX4 inhibition also improved the locomotor activity significantly and reduced the fatigue level by 22%. Our data overall demonstrate that the optimized antisense approach can contribute to future development of a therapeutic strategy for FSHD.

## Introduction

Facioscapulohumeral Muscular Dystrophy (FSHD) is a rare autosomal dominant genetic disorder with an estimated prevalence of 1:20,000 (1). The disease is characterized by asymmetric atrophy and weakness of the muscles of the face, shoulders and upper arms, which is extended to the trunk and lower limbs (2). Despite being the third most common muscular dystrophy, there is no existing disease-modifying treatment available for FSHD. This is partly because the complex mechanism underlying the disease has not been fully elucidated, although many candidate genes for FSHD have been identified (3–8). Of these, aberrant expression of the double homeobox 4 (*DUX4*) retrogene in skeletal muscle has been suggested in numerous studies to be predominantly involved in the pathogenesis of FSHD (9–14).

In healthy individuals, the subtelomeric region of chromosome 4q35 has 11-100 D4Z4 macrosatellite repeats, with each D4Z4 unit containing a copy of *DUX4* that encodes for a transcription factor, but is silenced in adult somatic tissues, including muscle (15). In most FSHD patients, contraction of the D4Z4 array to 1-10 repeats, associated with hypomethylation of the region, allows transcription of *DUX4* from the terminal D4Z4 repeat (15, 16). However, for the polyadenylation and stabilization of the *DUX4* transcript, and ultimately expression of the normally repressed DUX4 protein, the presence of a specific disease-permissive pLAM1 polyadenylation signal distal to the last D4Z4 unit is required (17). Inhibiting the activity of this structure leads to *DUX4* mRNA degradation through nonsense-mediated decay (18) and subsequently reduces protein synthesis. This has been achieved through the use of small interfering RNA (19), small hairpin RNA (20), microRNA (21), or antisense oligonucleotides (AONs) (22–24). Among these, AONs have several outstanding advantages. They can be administered without requirement of a viral vector, thereby avoiding triggering of an immune response and having a transient mode of action (25). This furthermore allows flexibility in dosage and frequency of dosing. In addition, AONs can be conjugated to a cell-penetrating moiety to enhance therapeutic efficiencies (26). Importantly, antisense therapy has emerged as a viable clinical path since four AONs received conditional FDA approval for use in subsets of patients with Duchenne muscular dystrophy (EXONDYS 51®, VYONDYS 53® and VILTEPSO®) or spinal muscular atrophy (SPINRAZA®).

We and others have reported that AON strategies were effective in downregulating *DUX4* expression in FSHD-derived myoblast cultures (19, 22), in a xenograft mouse model carrying muscle biopsies from FSHD patients (27) and as a local intramuscular treatment in a recently developed FLExDUX4 mouse model of FSHD (24, 28). Although these findings are promising, there remains lack of evidence of a systemic antisense effect; this is clinically important since a treatment for FSHD will need to suppress *DUX4* expression in a large number of skeletal muscles. Furthermore, expression of *DUX4* locally in the xenografted mice or at very low levels as seen in the FLExDUX4 model is not sufficient enough to recapitulate the phenotypes observed in FSHD patients (27, 29).

Following our promising *in vitro* finding (22), we have designed several AONs targeting both key elements in the 3’UTR of *DUX4* mRNA, the polyadenylation signal and the cleavage site. We initially investigated the antisense effect in FSHD-derived myoblasts and present here the enhanced efficacy of new sequences in downregulating expression of *DUX4* and its downstream targets to levels detectable in healthy isogenic myoblasts. We further studied the systemic therapeutic benefit of the best performing AON in the tamoxifen-inducible Cre-driver FLExDUX4 double-transgenic mouse model, where *DUX4* is inducibly expressed to a pathogenic level (30). We observed over 50% reduction in mRNA quantities of *DUX4* and downstream genes that led to significant amelioration in the muscle atrophy, muscle strength and bodywide activity of treated mice. Our novel findings suggest therapeutic potential of systemic delivery of the optimized antisense strategy as a treatment for FSHD.

## Results

### Newly designed antisense oligonucleotides significantly knock-down *DUX4* expression and ameliorate DUX4 pathology *in vitro*

We have previously shown that an antisense oligonucleotide (AON) with a phosphorodiamidate morpholino oligomer (PMO) backbone that targets the cleavage site (CS) of *DUX4* mRNA, PMO CS3, inhibited *DUX4* expression in FSHD myoblast cell cultures by over 50% (22). Another AON targeting the polyadenylation signal (PAS), located 20-30 nucleotides upstream of the *DUX4* CS was also effective in downregulating *DUX4* mRNA level by ∼40% (22, 27). Aiming to enhance the antisense efficacy against *DUX4*, we have designed four AONs that target both the PAS and CS, named PACS 1-4. Here, we assessed *DUX4* inhibitory effect of newly designed AONs relative to PMO CS3, considered as the positive PMO control. A PMO targeting the *HBB* mutation that causes β thalassemia was employed as a negative PMO control (PMO SCR). Details of PMOs used are shown in **Table S1**.

*In vitro* PMO screening was performed in FSHD-derived immortalized myoblast cells that have been characterized previously (31). The A5 clone containing 3 D4Z4 units represents patient cells whilst the A10 clone containing 13 D4Z4 units represents healthy control. We induced differentiation to FSHD immortalized myoblasts for 2 days then treated A5 cells with 10 μM of each PMO for 2 additional days. A5 and A10 cells receiving only the transfection reagent, Endo-Porter, were considered as untreated negative and positive controls, respectively (*n* = 3 per cell group). As shown in **Figure 1**, PMO SCR did not provide any inhibitory effect on any examined genes. Instead, all PACS PMOs were effective in downregulating mRNA levels of *DUX4* and three examined genes, which have been identified to be predominantly up-regulated in FSHD cells and muscle biopsies (32, 33), by 80-90% for *DUX4*, 65-85% for *PRAMEF2*, 53-68% for *TRIM43*, and 57-81% for *ZSCAN4* **(Figures 1a-d)**. *DUX4* expression following treatment with PMOs PACS1 and PACS2 remained at half of the level seen with PMO CS3 treatment, but was still significantly higher than the healthy level (*p*=0.0008 and 0.0035, respectively). PMOs PACS3 and PACS4 reduced *DUX4, PRAMEF2* and *ZCSAN4* expression to the level of A10 positive control, while the levels in PMO CS3 treatment remained significantly higher than the healthy values, specifically 6-fold for *DUX4* (*p<*0.0001), 25-fold for *PRAMEF2* (*p*=0.0192) and 13-fold for *ZSCAN4* (*p*=0.002).

**Figure 1:**
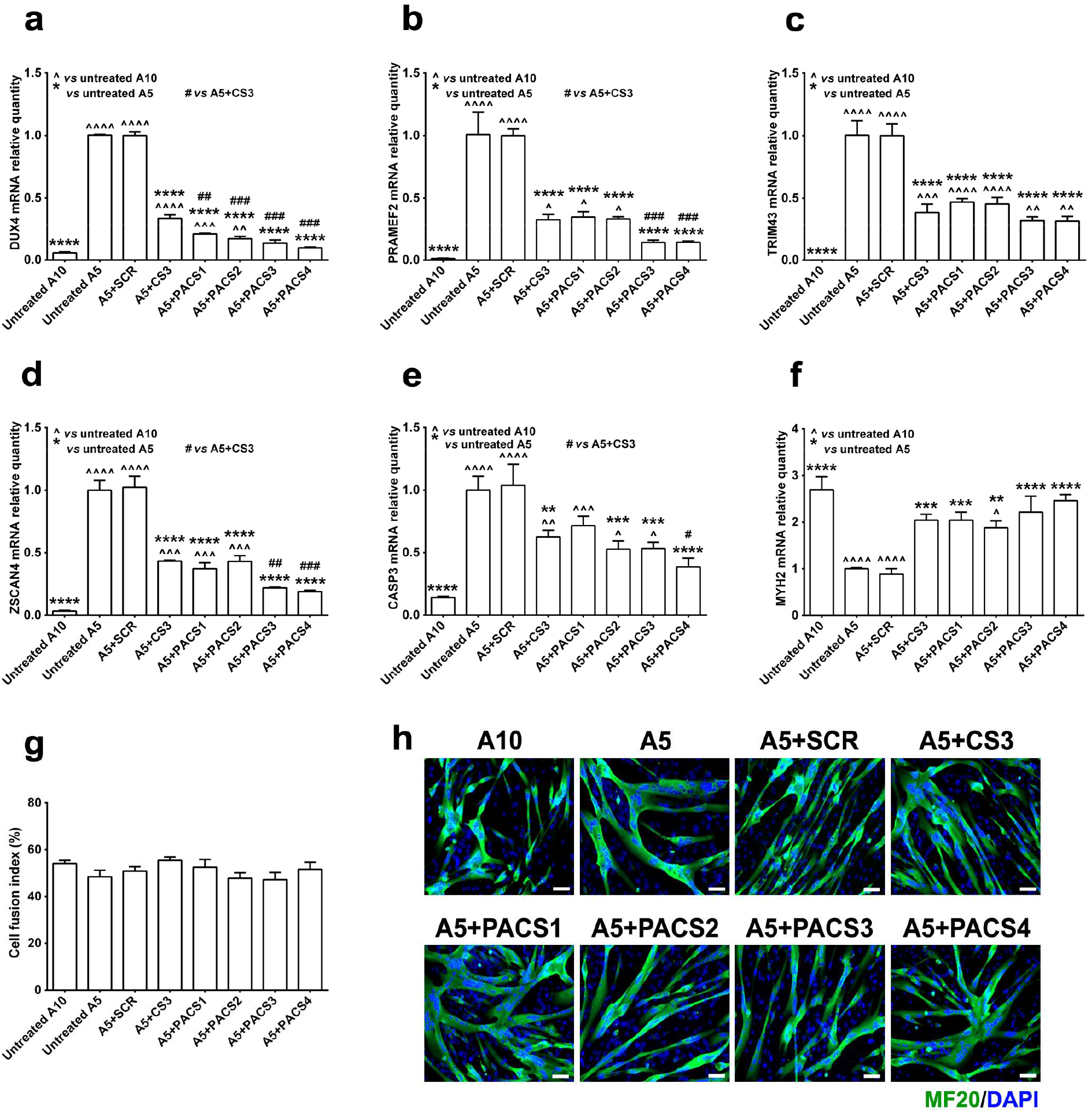
PMOs efficiently inhibit expression of *DUX4* and downstream targets in FSHD immortalized myoblast cell cultures. FSHD immortalized A5 myoblasts were differentiated for 2 days before the cells were treated with 10 μM PMOs through Endo-Porter-mediated transfection. Immortalized A10 or A5 cells receiving only Endo-Porter reagent were considered as untreated positive or negative control, respectively. Total RNA was extracted 2 days after PMO treatment. RT-qPCR quantification for *DUX4* **(a)** and its targets: *PRAMEF2* **(b)**, *TRIM43* **(c)**, *ZSCAN4* **(d)**, as well as markers of cell apoptosis *CASP3* **(e)** and cell differentiation *MYH2* **(f)** are shown, relative to corresponding *B2M* expression. Cells in a parallel study were immunostained for all myosin isoforms using MF20 antibody. Cell fusion indexes were evaluated as the number of nuclei in MF20-positive myotubes containing ≥ 3 nuclei and expressed as the percentage of the total nuclei number in the image field **(g)**. Representative cell images are displayed at magnification of x100, scale bar = 100 μm, MF20 (green), DAPI (blue) **(h)**. Statistical comparison **(a-g)** was by one-way ANOVA, followed by Tukey’s multiple comparisons test, and was against untreated A10 or A5 cell group, or A5 treated with PMO CS3 (considered as positive PMO control). Data are shown as mean ± S.E.M., *n* = 3, \**p* < 0.05, \*\**p* < 0.01, \*\*\**p* < 0.001, \*\*\*\**p* < 0.0001.

Since inappropriate expression of *DUX4* and its downstream genes has been reported to induce cell death and impair myoblast differentiation (34), we further assessed the mRNA level of *CASP3*, which is elevated in apoptotic cells, and *MYH2*, a marker of differentiated myoblasts. All PMOs, except PACS1, significantly reduced *CASP3* mRNA expression by 40-63%, relative to untreated A5. PACS4 provided the highest downregulation efficiency of 63% (*p*<0.0001) that was comparable to the level seen in the untreated A10 control (*p*=0.3642) and equivalent to 38% more inhibitory than CS3 treatment (*p*=0.0378) **(Figure 1e)**. Assessment of *MYH2* expression indicated that all DUX4-targeting PMOs significantly improved the mRNA level by 2-3-fold of the value in untreated A5, which was normalized to the healthy level of A10 control **(Figure 1f)**. However, the cell fusion index was comparable among all cell groups (*p*=0.2408), in consistency with previous findings (22, 31) **(Figure 1g)**. Representative images of cells immunostained for a myosin marker are shown in **Figure 1h**.

We also treated FSHD myoblasts with all PMOs at 1 μM. At this lower dose, newly designed PMOs remained effective in inhibiting expression of *DUX4* by 27-44%, *PRAMEF2* by 47-65%, *TRIM43* by 26-51%, and *ZSCAN4* by 27-38%, although they were not significantly more efficient than PMO CS3 treatment **(Figures S1a-d)**. PMO PACS4 continued to be the best performing candidate and was the only PMO that could reduce *CASP3* expression to the level seen in the A10 positive control (*p*=0.6570), **(Figure S1e)**. Treatment with PACS4 also increased significantly the *MYH2* mRNA level relative to untreated A5 myoblasts by 2-fold (*p* = 0.0087), or by 1.5-fold as compared to CS3-treated cells (*p* = 0.0167) **(Figure S1f)**.

Based on these *in vitro* data, PMO PACS4 was significantly more effective than PMO CS3, the best candidate identified previously, in inhibiting expression of *DUX4* and other FSHD-related genes and therefore was selected for the animal study.

### Systemic antisense treatment efficiently improves the mass of several skeletal muscles in a tamoxifen-induced FSHD mouse model

To study the *in vivo* antisense effect, we employed tamoxifen (TMX)-inducible Cre-driver FLExDUX4 double transgenic mice, named MCM-D4. Extensive characterization by the Jones Lab indicated that TMX-mediated induction can be either very mild or too severe depending on the dose regimen (29, 30). Hence, to tailor the model for testing *in vivo* antisense treatment, we initially optimized the dose regimen of TMX for inducing *DUX4* expression. Sixteen-week-old MCM-D4 males were injected with either a single dose of 5 mg/kg TMX (*n*=5) or 2.5 mg/kg/biweekly TMX (*n*=5) via intraperitoneal (IP) administration. Age-matched HSA-MCM mice receiving volume-matched corn oil were considered as wild-type (WT) control (*n*=5). The body weight recorded weekly **(Figure S2a)** indicated a complete recovery of 100.9% start weight at week 4 following 5 mg/kg TMX induction, similar to WT level of 104.5% (*p*=0.0668), and consistent with the characterization by the Jones Lab (29, 30). Although the same overall TMX dose was used, biweekly administration of 2.5 mg/kg TMX led to chronic weight loss and reached 12.5% (*p*<0.0001) of the level seen in mice receiving a single dose at the end of the experiment. Moreover, the mass of tibialis anterior (TA) correlated with the effect of TMX dosage, with 26% (*p*<0.0001) more muscle wasting observed following biweekly TMX injection than the single administration **(Figure S2b)**. *In situ* TA force measurement additionally demonstrated 30% (1015 ± 19.13 mN, *p*<0.0001) or 65% (509 ± 87.24 mN, *p*<0.0001) drop of WT force (1438 ± 50.21 mN) in mice injected with 5 mg/kg or 2.5 mg/kg/biweekly TMX, respectively **(Figure S2c)**. These data together suggested that the biweekly administration of 2.5 mg/kg TMX successfully generated a model with progressive DUX4-mediated muscle atrophy; therefore, this dose regimen was used to investigate our antisense approach *in vivo*.

Following promising results from *in vitro* screening and aiming to enhance cell penetration in murine muscle (26), we conjugated PMO PACS4 with an octaguanidine dendrimer chemistry, named Vivo-PMO PACS4. MCM-D4 males (16-week-old) were injected with 2.5 mg/kg TMX on days 0 and 14. Age-matched HSA-MCM males receiving the same TMX dosage were considered as WT controls (*n*=4). MCM-D4 mice were further IP injected with 10 mg/kg of Vivo-PMO PACS4 (*n*=5), or 10 mg/kg of Vivo-PMO SCR (*n*=4), considered as negative control, on days 2, 8, 16, and 22 whilst HSA-MCM mice received volume-matched saline. Body weight recorded during the study, just prior to injections or functional tests, and normalized to the initial weight **(Figure 2a)**, displayed significant weight loss by ∼6% in MCM-D4 versus WT mice from day 11 (*p*=0.0023) that continued to drop by 12% (in PACS4, *p*<0.0001) or 16% (in SCR, *p*<0.0001) at the end of experiment. In comparison to Vivo-PMO SCR, treatment with PACS4 significantly improved the body weight on day 22 by 4% (*p*=0.0109) and remained effective for the course of the treatment (*p*=0.0027). In coherence with body weight improvement, PACS4 administration significantly improved the mass of three examined muscles, including the gastrocnemius (GAS) from 4.37 ± 0.06 to 5.03 ± 0.04 mg/g (*p*<0.0001), quadriceps (QUAD) from 4.72 ± 0.14 to 5.44 ± 0.08 mg/g (*p*=0.0109), and tibialis anterior (TA) from 1.35 ± 0.02 to 1.42 ± 0.01 mg/g (*p*=0.0189) that were normalized to the corresponding body weight **(Figures 2b-d)**. Thereby, PACS4 treatment increased the mass of these muscles by up to 15% of the values of SCR-treated tissue.

**Figure 2:**
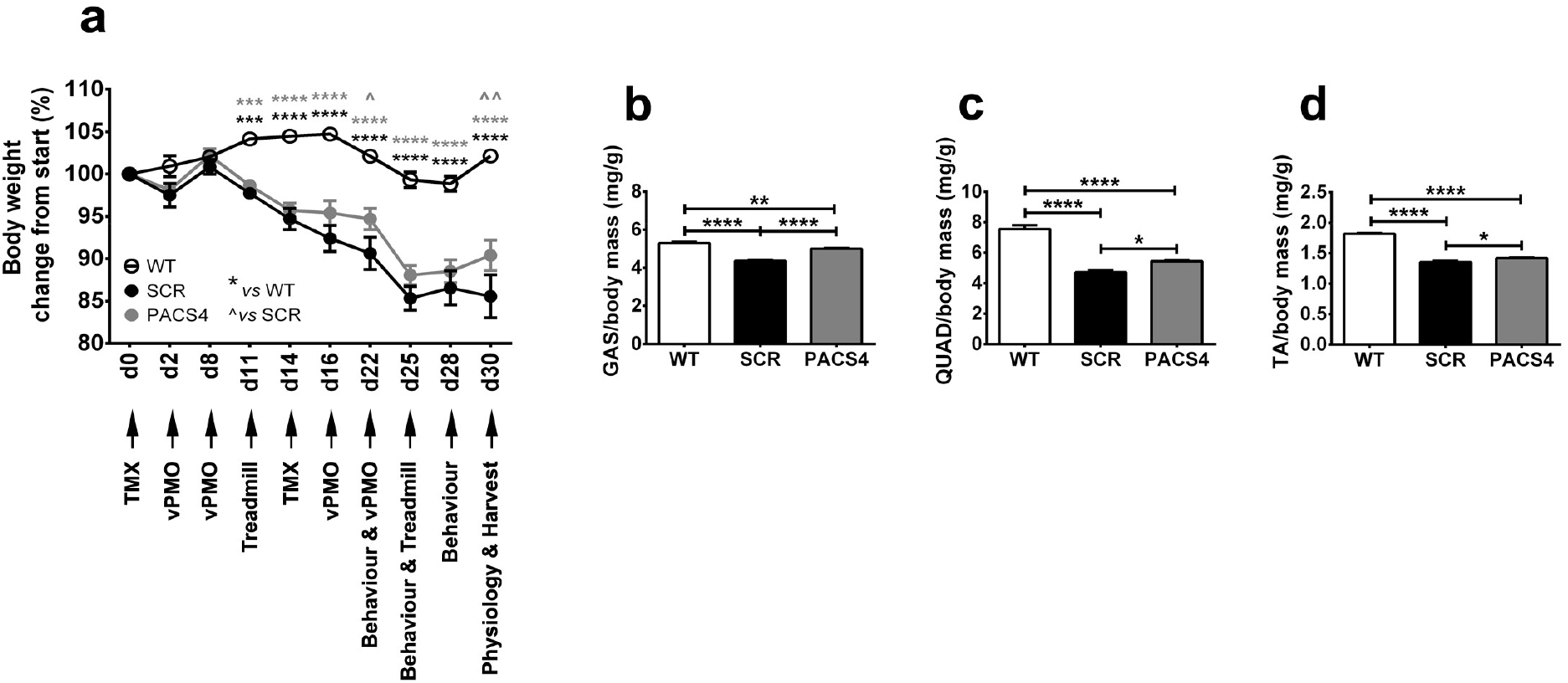
*In vivo* experimental plan and effect of Vivo-PMO PACS4 treatment on body and muscle mass. The effect of antisense therapy was studied in 16-week-old mice. Details of the experiment, including timepoints for functional tests, and changes in body weight of mice are presented **(a)**. All mice received 2.5 mg/kg/biweekly tamoxifen (TMX) by intraperitoneal (IP) injection. MCM-D4 mice were further injected with 10 mg/kg/week of Vivo-PMO PACS4 (n = 5), or 10 mg/kg/week of Vivo-PMO SCR (n = 4), considered as a negative control. HSA-MCM mice receiving volume-matched saline, instead of Vivo-PMO, were considered as wild-type (WT) controls (n = 4). Animals were sacrificed 1 week after the last Vivo-PMO injection. Muscle mass of the gastrocnemius - GAS **(b)**, quadriceps - QUAD **(c)**, and tibialis anterior - TA **(d)** normalized to the corresponding body weight is displayed. Data are shown as mean ± S.E.M. Statistical analysis was by two-way **(a)** or one-way ANOVA **(b-f)** followed by Tukey’s multiple comparisons test, \**p* < 0.05, \*\**p* < 0.01, \*\*\**p* < 0.001, \*\*\*\**p* < 0.0001.

### Systemic administration of Vivo-PMO PACS4 improves locomotor behavior, whole body function and TA muscle strength

After 4 weekly Vivo-PMO injections, we assessed the locomotor behavior of the animals using open-field activity cage monitors. MCM-D4 mice appeared less active than WT controls in 16 of 22 parameters examined **(Table S2)**. However, following PACS4 treatment MCM-D4 mice displayed a significant improvement in 11 parameters relative to SCR-treated mice. Of importance, the total active time, total rearing time and travelled distance were all increased, from 215.4 ± 41.3 sec to 584.5 ± 64.0 sec (*p*<0.0001), 59.9 ± 14.3 sec to 182.6 ± 18.1 sec (*p*=0.0238), 14.6 ± 3.1 m to 31.8 ± 3.6 m (*p*=0.0116), respectively **(Figures 3a-c)**. We also evaluated the effect of antisense treatment on whole body function using treadmill exhaustion test. Mice were allowed to run on a treadmill until they were unable to move from the stopper for 10 sec. The total running times were then calculated as the percentage of the baseline time recorded prior to the beginning of the experiment and presented in **Figure 3d**, indicating that the fatigue level was significantly higher in SCR-injected mice than in WT controls just after the first TMX injection by 37% at week 2 (*p*=0.0013) and 62% at week 4 (*p*<0.0001). In contrast, mice receiving PACS4 treatment were 27% (*p*=0.0118) and 22% (*p*=0.0249) less exhausted than SCR-treated mice, respectively and only exhibited significant fatigue relative to WT at week 4, by 40% (*p*=0.0007).

**Figure 3:**
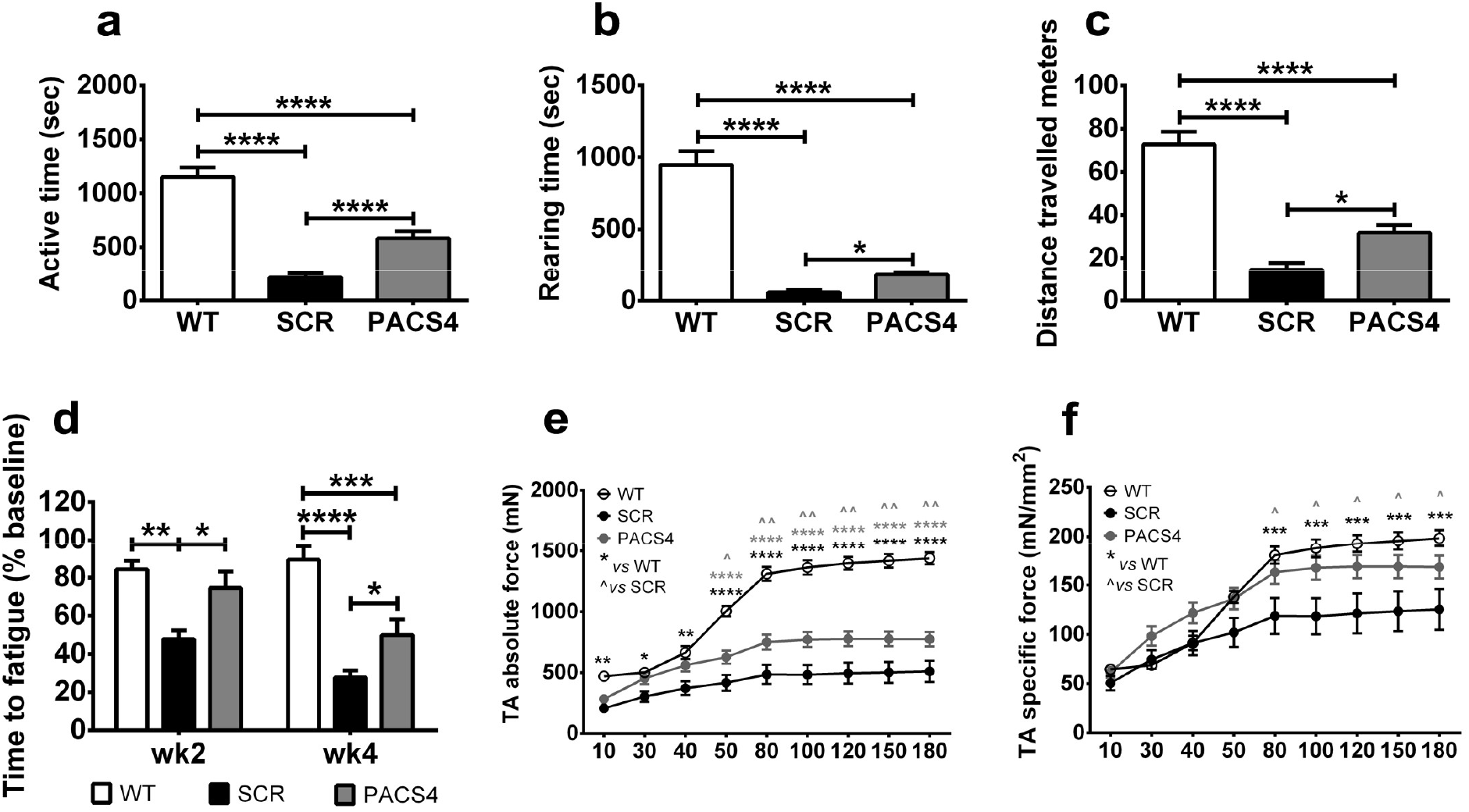
Systemic antisense therapy improves animal behavior and muscle function. Mouse open-field behavior was assessed using locomotor activity monitors. Representative parameters of the animal behavior are shown in **(a-c)** while details of all parameters are presented in *Table S2*. Antisense efficacies were further evaluated via treadmill exhaustion tests, ∼1 week prior to the first TMX injection (baseline), then at weeks 2 and 4. Total running time on treadmill was recorded and expressed as the time to fatigue as the percentage of baseline time **(d)**. Following 4 weekly Vivo-PMO administration, mice were put under terminal anesthesia and *in situ* TA absolute force was measured **(e)**. Specific muscle force is displayed as a ratio of the absolute force and the TA cross-sectional area **(f)**. Data are shown as mean ± S.E.M, *n* = 4-5. Statistical analysis was by one-way ANOVA **(a-c)** or two-way ANOVA **(d-f)**, followed by Tukey’s multiple comparisons test, \**p* < 0.05, \*\**p* < 0.01, \*\*\**p* < 0.001, \*\*\*\**p* < 0.0001.

Additional *in situ* TA force measurement demonstrated obvious muscle weakness in MCM-D4 mice receiving Vivo-PMO SCR, compared to WT mice **(Figures 3e, f)**. Treatment with PACS4 significantly improved the absolute muscle force from 509.4 ± 87.2 mN to 773.8 ± 59.4 mN (*p*=0.0049 at 180 Hz), relative to SCR-treated mice. Importantly, mice receiving PACS4 exhibited specific muscle force comparable to WT mice, and that was significantly stronger than SCR-treated mice (125.4 ± 20.7 mN/mm^2^ versus 168.4 ± 11.6 mN/mm^2^, *p*=0.0351 at 180 Hz).

### Vivo-PMO PACS4 robustly inhibits mRNA expression of *DUX4* and downstream targets

Since DUX4-induced pathology of TA muscle is a clinical characteristic of FSHD (2, 35, 36), the muscle has been extensively studied in numerous pre-clinical FSHD research (27, 30, 37–40). Hence, we focused our investigation on the effect of Vivo-PMO PACS4 treatment in TA muscle. To verify the antisense efficacy on mRNA expression, we carried out RT-qPCR quantification for *DUX4* and three downstream genes that have been proven to be activated in FSHD, including *Frg1, Trim43* and *Wfdc3* (3, 30, 41) **(Figures 4a-d)**. As predicted, the mRNA levels of all genes were greatly elevated in SCR group, relative to the WT values. Treatment with PACS4 reduced mRNA quantities of the examined genes by about half of the SCR levels (*p*=0.0453, 0.0007, 0.0146, 0.0063, respectively). Further evaluation of the Vivo-PMO effect on muscle apoptosis and consequent muscle regeneration by assessing the mRNA levels of *Casp3, Myh3*, and *Pax7* **(Figures 4e-g)** demonstrated that gene expression following SCR administration was upregulated by 7-fold (*p*<0.0001), 96-fold (*p*=0.0052), and 4-fold (*p*=0.0075), as compared to WT levels, respectively. PACS4 treatment significantly downregulated expression relative to the SCR values by ∼50% (*p*=0.0071, 0.0093 and 0.0461, respectively).

**Figure 4:**
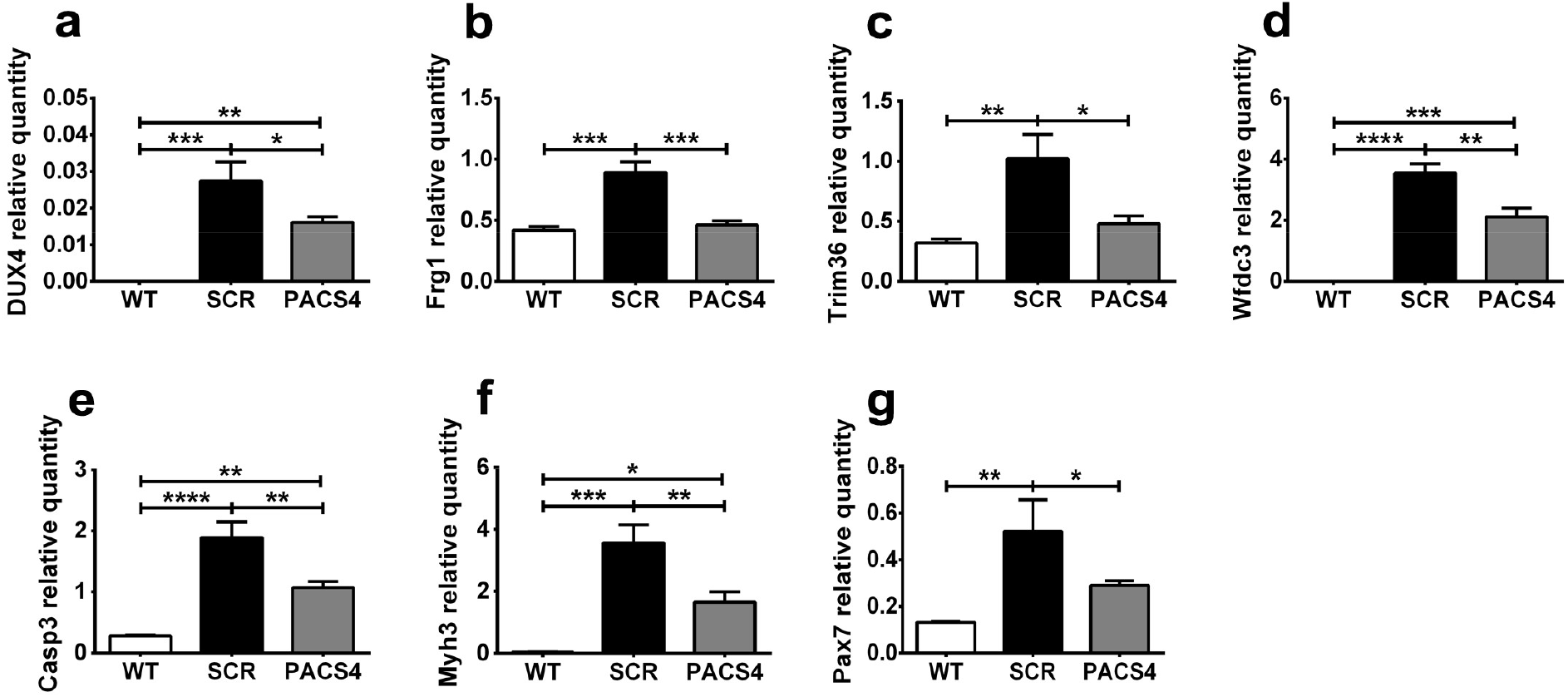
Vivo-PMO PACS4 treatment downregulates expression of *DUX4* and relevant genes. Following TMX-induced *DUX4* expression and 4 weekly treatment with Vivo-PMOs, changes at mRNA levels in TA muscle was examined by RT-qPCR for *DUX4* **(a)**, *Frg1* **(b)**, *Trim36* **(c)**, *Wfdc3* **(d)**, *Casp3* **(e)**, *Myh3* **(f)** and *Pax7* **(g)**, relative to corresponding *Gapdh* expression. Data are shown as means ± S.E.M; *n* = 4-5. Statistical comparison was by one-way ANOVA followed by Tukey’s *post-hoc* test; \**p* < 0.05, \*\**p* < 0.01, \*\*\**p* < 0.001, \*\*\*\**p* < 0.0001.

### PACS4-mediated DUX4 inhibition greatly improves muscle histopathology

To assess the effect of the treatment with Vivo-PMO PAS4 on muscle histopathology, TA muscle sections were analyzed. Muscle sections were immunostained for laminin to assist the identification of the myofiber sarcolemma for subsequent analyses of the cross-sectional area (CSA), the number of total fibers and number of centrally nucleated fibers (CNFs). We observed a decrease in the CSA of MCM-D4 muscle, and that PACS4 treatment significantly rescued the muscle atrophy relative to SCR-injected mice by 12%, from 5.3 ± 0.2 to 6.7 ± 0.2 mm^2^ (*p*=0.0268) **(Figure 5a)**. The number of TA myofibers/mm^2^ in both MCM-D4 groups was significantly higher (*p*<0.0001) than the WT value **(Figure 5b)**. Together with elevated *Casp3* and *Myh3* mRNA expression, this suggested myofibers turning-over due to TMX-induced DUX4 expression. An increase in the number of myofibers expressing embryonic MyHC (eMyHC), a marker of muscle regeneration confirmed this hypothesis; although, the eMyHC number was significantly less in TA injected with PACS4 (11.6 ± 1.9) than with SCR (16.8 ± 1.8), *p*=0.0401 **(Figure 5c)**, in consistency with mRNA quantification for *Myh3*.

**Figure 5:**
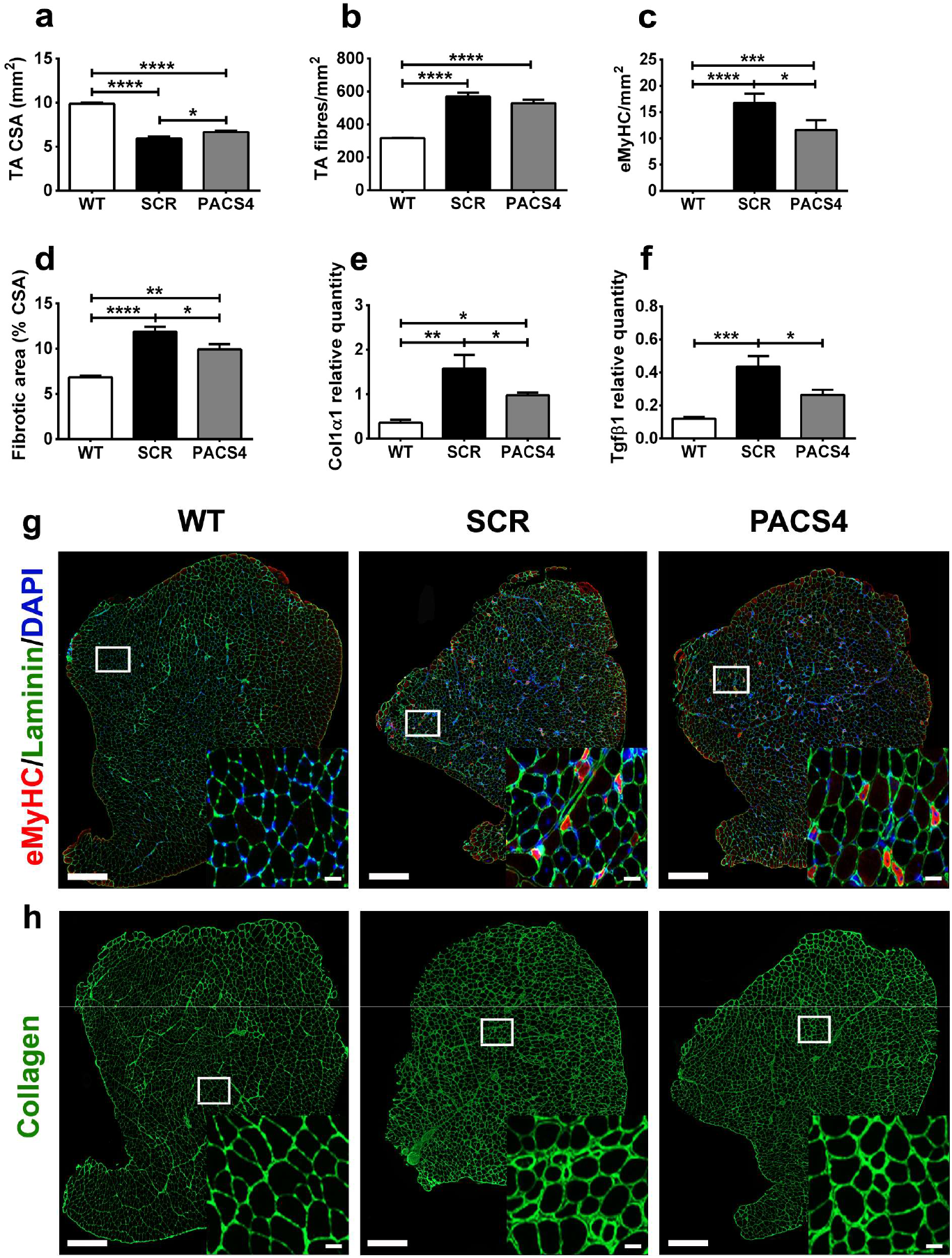
Effect of Vivo-PMO PACS4 therapy on muscle histopathology. Frozen TA muscle sections were stained for laminin, embryonic myosin heavy chain (eMyHC), and DAPI. The cross-sectional area (CSA) of the entire muscle section was automatically scored by MuscleJ **(a)**. Laminin staining was used for identifying the fiber perimeter. An average of 3200 myofibers/TA was examined and displayed as the total number of myofibers **(b)** or the number of myofibers positive with eMyHC staining **(c)**, per mm2 of the CSA. Fibrotic area in TA muscle was semi-automatically evaluated and expressed as percentage of the area positive for collagen of the muscle CSA **(d)**. mRNA expression of gene indicative for fibrotic response, *Col1α1* **(e)** and *Tgfβ1* **(f)**, was quantified by RT-qPCR. Statistical comparison was by one-way ANOVA followed by Tukey’s *post-hoc* test **(a-f)**. Data are shown as means ± S.E.M; *n* = 4-5; \**p* < 0.05, \*\**p* < 0.01, \*\*\**p* < 0.001, \*\*\*\**p* < 0.0001. Representative images of the entire TA cross-sections co-stained with eMyHC (red), laminin (green), DAPI **(g)**, or single stained with collagen VI **(h)** are shown at magnification of x100, scale bar = 500 μm. Corresponding enlarged images at higher magnification are shown in the subsets, scale bar = 50 μm **(g, h)**.

Further examination on the level of excessive muscle fibrosis, based on immunostaining for a common marker of fibrosis, collagen VI (42, 43), revealed that *DUX4* expression led to increase in the fibrotic area of TA muscle, from 6.82 ± 0.21% (WT) to 11.89 ± 0.54% (SCR, *p*<0.0001). PACS4-treated muscle displayed 9.93 ± 0.57% fibrosis, significantly reduced by 17% of the value in SCR muscle (*p*=0.0403) **(Figure 5d)**. Since the level of fibrosis was upregulated in MCM-D4 muscle, we explored mRNA expression of genes indicative of a fibrotic response, *Col1α1* and *Tgfβ1* **(Figures 5e, f)**. As expected, *Col1α1* and *Tgfβ1* levels were higher in both SCR and PACS4 groups relative to the level in WT. However, PACS4 treatment significantly reduced expression relative to that seen in SCR muscle by 38% for *Col1α1* (*p*=0.0421) and 40% for *Tgfβ1* (*p*=0.0296), in agreement with the histological analysis. Representative TA images from immunostaining for eMyHC and collagen VI are shown in **Figures 5g, h**.

### Antisense treatment prevents a switch in myofiber types and improves myofiber atrophy

We initially assessed gene expression of four major myofiber types in mammalian skeletal muscles, including *Myh7* for slow-twitch MyHC I, *Myh2* for fatigue-resistant fast-twitch MyHC IIA, *Myh4* for fatigable fast-twitch MyHC IIB, and *Myh1* for fast-twitch MyHC IIX (fatigue resistance less than IIA but better than IIB) (44). Expression of *Myh7* was undetectable in all TA groups. *Myh2* and *Myh1* levels in MCM-D4 muscle were lower than WT values by 40% (*p*=0.0396) and 48% (*p*=0.0160) in SCR-treated muscle, and by 59% (*p*=0.0742) and 57% (*p*=0.0247) in PACS4-treated muscle, respectively **(Figures S4a, b)**. In contrast, *Myh4* expression was upregulated by 141% (*p*=0.0437) in SCR group, while treatment with PACS4 significantly reduced the level by 64% (*p*=0.0213), towards the WT value (*p*=0.4932) **(Figure S4c)**. According to this finding, we performed immunostaining for laminin, MyHC IIA and IIB on TA transverse sections; unstained myofibers were considered as type IIX. Results from automatic MuscleJ quantification indicated a shift in myofiber populations, in consistency with the qPCR analysis above. MCM-D4 muscle displayed a decrease in the percentage of types IIA and IIX, but an increase in type IIB, significantly as compared to the WT **(Figures 6a-c)**. PACS4 group had similar quantity of type IIA relative to the SCR group (2.88% *vs* 2.65%, *p*=0.8046). However, the percentage of MyHC IIX in PACS-treated TA was higher than in the SCR group (30.6% *vs* 24.7%, *p*=0.0489) while the level of type IIB was respectively lower by 8.5% (*p*=0.0362), towards the WT properties. Representative images of immunostained TA sections and of MuscleJ analysis are shown in **Figure 6j**.

**Figure 6:**
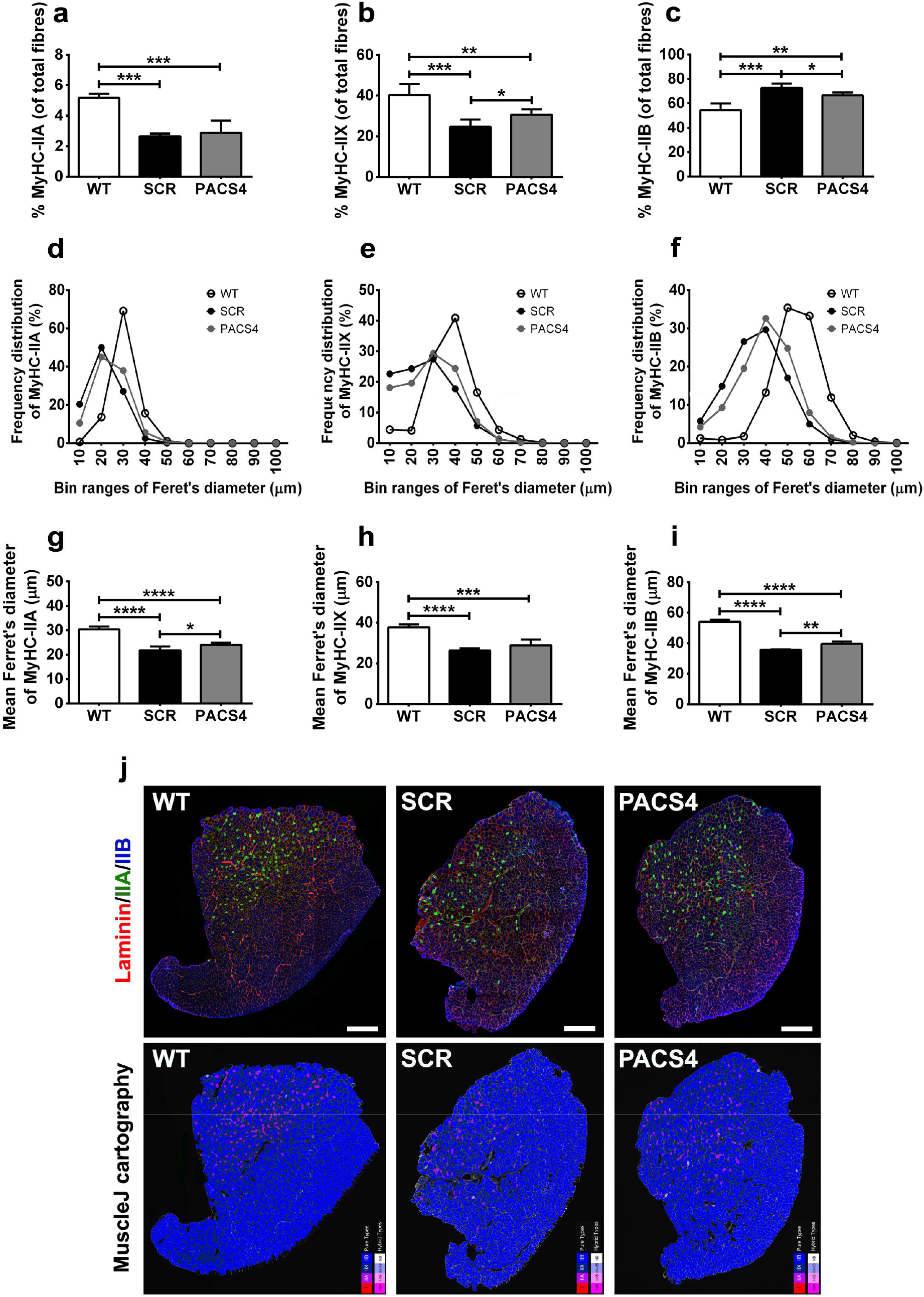
Vivo-PMO PACS4-mediated amelioration of muscle atrophy and myofiber type switching. Frozen TA muscle sections were immunostained for laminin (red), MyHC IIA (green), MyHC IIB (blue); unstained fibers were considered as MyHC IIX. The number of MyHC-positive fibers was automatically scored by MuscleJ and is expressed as the percentage of the total number of all myofibers within the entire muscle sections **(a-c)**. Laminin staining was used for identifying the fiber sarcolemma and subsequent analysis of the minimal Feret’s diameter of myofibers. Histograms of frequency distribution **(d-f)** and the mean of the diameter of each myofiber type **(g-i)** are presented. Data are shown as mean ± S.E.M., *n* = 4-5. Statistical comparison was by one-way ANOVA and Tukey’s *post-hoc* test **(a-c, g-i)**; \**p* < 0.05, \*\**p* < 0.01, \*\*\**p* < 0.001, \*\*\*\**p* < 0.0001. Representative images of the entire TA muscle sections are shown at magnification of x100, scale bar = 500 μm **(j)**. Corresponding cartographic images created by MuscleJ software are also displayed, with MyHC IIA, IIB and IIX fibers color-coded as purple, light blue and dark blue, respectively.

Evaluation of the minimal Feret’s diameter of myofibers indicated myofiber atrophy occurring in all myofiber types of MCM-D4 mice **(Figures 6d-i)**, in line with the decrease in the CSA presented above. Histograms of the frequency distribution of the fiber size demonstrated that the majority of WT MyHC IIA, IIX and IIB fibers was 30, 40 and 50 μm in diameter respectively whereas the Ferret’s diameter of corresponding MCM-D4 myofiber types peaked at 10-μm smaller values **(Figures 6d-f)**. Subsequent calculation of the mean fiber diameter clarified that the sizes of SCR-treated MyHC IIA, IIX and IIB fibers, relative to type-matched WT fibers, decreased from 30.4 ± 0.6 to 21.8 ± 0.8 μm (*p*<0.0001), 37.8 ± 0.7 to 26.3 ± 0.5 μm (*p*<0.0001) and 54.1 ± 0.6 to 35.7 ± 0.2 μm (*p*<0.0001), respectively. PACS4 did not alter the proportion of MyHC IIA, but significantly increased the myofiber diameter, compared to SCR treatment, from 21.8 ± 0.8 to 24.1 ± 0.4 μm (*p*=0.0424). Administration of PACS4 was also effective in improving the mean fiber size of MyHC IIB, relative to SCR values, from 35.7 ± 0.2 to 39.7 ± 0.7 μm (*p*=0.0014), **Figures 6g-i**.

Taken together, histological analyses provided evidence for DUX4-mediated muscle atrophy in the TA of MCM-D4 mice, in consistency with the reduction seen in the mass and strength of the muscle. Treatment with Vivo-PMO PACS4 efficiently ameliorated DUX4 histopathology by preventing myofiber atrophy and shifts in the myofiber type profile.

## Discussion

Despite increased understanding of genetic and epigenetic factors that contribute to FSHD, there is no treatment that can prevent or delay the disease progression. Clinical management involving physiotherapies, vision and hearing aids, orthopedic interventions, pain and fatigue management, or surgical scapular fixation, has shown some clinical benefit and improved the quality of life for FSHD patients (35, 45). Since aberrant expression of *DUX4* gene has been extensively reported as the main causative factor for FSHD (9–14, 19), and the stabilization of *DUX4* mRNA requires recognition of the polyadenylation signal (PAS) and the cleavage site (CS) in the 3’UTR, we developed antisense oligonucleotides (AONs) with phosphorodiamidate morpholino oligomer (PMO) chemistry that target both the PAS and the CS. We demonstrate in this study that newly designed PMOs improved the antisense efficacy of previous AONs targeting either the PAS or the CS (22, 27), inhibiting mRNA expression of *DUX4* and its downstream genes to levels seen in isogenic positive control cells. All new PMOs were significantly more effective at reducing *DUX4* expression compared to the PMO CS3 control. PMOs PACS 3 and particularly PACS 4 also provided greater inhibition than CS3 on DUX4 targets. These results indicate that further optimization of the AON sequences can improve the impact of the antisense therapy. AONs targeting other locations in the *DUX4* 3’UTR have been also proven efficacious in *in vitro DUX4* inhibition (24, 28). However, *in vivo* investigation of such AONs did not include any muscle functional or histopathological assessment; hence, such trials could not provide a complete pre-clinical examination of the therapeutic benefit of the approach (24, 28). Importantly, they have only been tested by local intramuscular injection, while any treatment for FSHD is crucially required to have a systemic effect.

Here, we show for the first time that systemic delivery of antisense therapeutics improves FSHD disease in a relevant mouse model. We demonstrated increase in the mass of the examined skeletal muscles, improved locomotor behavior, and amelioration in the fatigue level of treated mice. The benefit to the tibialis anterior (TA) muscle was highlighted by the antisense inhibitory effect on mRNA expression of *DUX4* and its downstream targets, and the consequent beneficial effects on markers of the apoptosis, regeneration and fibrosis of the muscle. Most importantly, therapeutic outcome was evidenced by enhancement in the *in situ* force generating capacity and amelioration in the atrophy of the examined tissue. Increase in muscle strength was likely at least partially due by antisense-mediated prevention in the shift of myofiber populations from MyHC IIX to IIB. In fact, in FSHD, MyHC IIB fibers have been proven to generate much less force and display less mitochondria than other fiber types (46, 47), and that contributed to the pathology observed in the *FRG1* expressing mouse model of FSHD (46). Therefore, we speculate that by suppressing the switch into MyHC IIB fibers our antisense treatment could provide additional beneficial effects in the mitochondrial content and function, and consequently in energy metabolism; this will need to be investigated in future work.

Despite these promising findings, our approach needs to overcome two major challenges to support its translation into clinical setting for FSHD. The first obstacle is the efficiency of AON uptake into myofibers and its biostability. PMO has good life time in skeletal muscle and shows no serious toxic concern in treatment for Duchenne muscular dystrophy (DMD). Nevertheless, as a charge-neutral chemistry, PMO displays poor cellular uptake which means it does not easily penetrate into muscles of FSHD patients. Conjugating PMO with a cell-penetrating moiety, for instance the Vivo-PMO used here, has improved cell-penetrating efficacy (48, 49). As demonstrated, the antisense benefit of Vivo-PMO PACS4 can be achieved systemically in non-leaky muscles of the murine model used. Vivo-PMO further has enhanced stability due to its arginine-rich component (48), although this modification may cause unwanted effects (50). Therefore, to obtain the best therapeutic benefit while minimizing the side-effects, it is important to monitor the sensitivity of the organisms to be treated and adjust the dosage over the treatment course, as it has been done successfully by many groups including ourselves (26, 49, 51, 52). PMO can be also conjugated with alternative cell-penetrating peptides (26, 53–55), or be delivered via nanoparticles (56) or even packaged into adeno-associated viral vectors (57) that have been shown to improve cellular uptake in many pre-clinical studies; even so, clinical safety of these alternative chemistries and delivery mechanisms requires further meticulous investigation.

The second challenge for translational development of antisense therapies, as well as of other approaches for FSHD, is the lack of a standard animal model of the disease. There are four animal models of FSHD available, including the D4Z4-2.5 (41), iDUX4pA (58), FLExDUX4 (29), and Rosa26-DUX4 (38) mice. Many efforts have been additionally made to improve these models to recapitulate better the complicated pathophysiological mechanism of DUX4 action and the phenotypic features seen in FSHD patients (30, 59), but further optimization remains essential. For example, the specific TMX dose regimen we developed in this study for the Cre-driver FLExDUX4 mice can overcome the innate muscle recovery of the previous model introduced by Jones et al (30); however, for studying the long-term potential of therapeutic applications for FSHD, extensive optimization is needed to obtain a more chronic model while minimizing the risk of lethality caused by DUX4 accumulation over time. Further, as skeletal muscles are not equally affected in these models, consistently as seen in patients with FSHD (31, 60), and DUX4 has been detected in biopsies of unaffected individuals (61), more research is needed to determine whether DUX4 has function in other tissues and the level of DUX4 suppression needed to obtain clinical therapeutic outcomes.

In summary, the present study provides substantial evidence that systemic *in vivo* treatment with an improved antisense design results in reduction in mRNA quantities of *DUX4* and target genes that leads to amelioration in the muscle function, muscle histopathology, and locomotor activities of treated mice. Our data overall demonstrate that the optimal antisense approach can contribute to future development of a therapeutic strategy for FSHD.

## Materials and Methods

### PMOs and Vivo-PMOs

Phosphorodiamidate morpholino oligomers (PMOs) and octaguanidine dendrimer-conjugated PMOs (Vivo-PMOs) were purchased from GeneTools (Oregon, USA). To allow conjugation to octaguanidine dendrimer, 28-mer version of the optimized PMO was used in the *in vivo* work. PMOs and Vivo-PMOs were dissolved in sterile ddH_2_O and were further diluted to desired concentrations in cell culture medium (*in vitro* work) or in sterile 0.9% saline (Sigma, UK) immediately prior to injection into mice. Sequences of the PMOs and Vivo-PMOs are listed in *Table S1*.

### Cell cultures and PMO transfection

FSHD immortalized myoblast cells that have been characterized previously (31) were kindly provided by Dr Vincent Mouly, Institute of Myology, France. The A5 clone containing 3 D4Z4 units was considered as being contracted, whilst the A10 clone containing 13 D4Z4 units was considered as being non-contracted and used as a positive control. Cells were maintained in proliferation medium composed of 64% (v/v) high glucose DMEM (Gibco, UK), 16% (v/v) Medium 199 (Gibco, UK), 20% (v/v) foetal bovine serum (FBS, Gibco, UK), 50 μg/ml gentamicin (Sigma, UK), 0.2 μg/ml dexamethasone (Sigma, UK), 0.5 ng/ml human basic fibroblast growth factor (Sigma, UK), 5 ng/ml human recombinant epidermal growth factor (Sigma, UK) and 25 μg/ml fetuin from FBS (Sigma, UK). Cell differentiation was induced when cells reached around 90% confluence by replacing the proliferation medium with 99% (v/v) high glucose DMEM (Gibco, UK), 1% (v/v) horse serum (Gibco, UK) and 10 μg/ml human insulin-transferrin-sodium selenite media supplement (Sigma, UK). To study the antisense inhibitory effect, myoblasts were differentiated for 2 days and treated with 1 or 10 μM PMOs, *n* = 3 per cell group. Transfection was facilitated by 6 μM Endo-Porter (GeneTools, Oregon, USA). RNA extraction was conducted after 2 additional days.

### RT-qPCR quantification for *DUX4* and relevant genes

Total RNA from cultured cells was extracted using RNeasy kit (QIAgen, UK) while RNA from murine muscles (as described in the animal study below) was extracted using RNeasy Fibrous Tissue kit (QIAgen, UK), following the manufacturer’s instructions. Tissue homogenisation was performed in the lysis buffer provided with the kit, at 25 Hz for 2-4 min, on a TissueLyser II (QIAgen, UK). RNA was quantified on a ND-1000 NanoDrop spectrophotometer (Thermo Scientific, UK). One microgram RNA was reverse transcribed using QuantiTect reverse transcription kit (QIAgen, UK). Ten nanograms of diluted cDNA in qPCR water (Roche, UK) were then amplified using LightCycler480 SYBR Green Master I kit (Roche, UK), according to the manufacturer’s instructions; samples were prepared in triplicates. Reactions were run on LightCycler480 System, initialized at 95°C for 5 min, followed by 45 cycles at 95°C for 15 sec, 58°-62°C for 15 sec, 72°C for 15 sec. Relative quantification for *DUX4* or its relevant genes was performed against corresponding housekeeping genes, *B2M* or *Gapdh*. Primers were purchased from Integrated DNA Technologies (Belgium) and are detailed in *Table S3*.

### Immunocytochemistry for quantifying cell fusion index

FSHD immortalized myoblast cells were seeded in 6-well plates pre-coated with extracellular matrix gel from Engelbreth-Holm-Swarm murine sarcoma (Sigma, UK). Following transfection with PMOs, the culture medium (as described above) was removed on day 4 of cell differentiation. Cells were rinsed in ice-cold 1x PBS (Sigma, UK), fixed in ice-cold 4% (w/v) paraformaldehyde (Sigma, UK) for 10 min and permeabilised in 1x PBST (0.25% (v/v) Triton X-100, 1x PBS, Sigma, UK) for 10 min. Cells were blocked in 1% (w/v) bovine serum albumin, BSA (Sigma, UK), 10% (v/v) goat serum (Sigma, UK), 1x PBST for 1 hr. Incubation with mouse anti-myosin heavy chain (MF20) primary antibody (1:100, DSHB, Iowa, USA) was performed at 4°C overnight, followed by incubation with goat anti-mouse AlexaFluor488 secondary antibody (1:500, Life Biotechnologies, UK) for 1 hr at room temperature. MF20 antibody was generated and deposited to DSHB by Fischman, D.A. (62). Nuclei were further stained with 1 μg/ml 4′,6-diamidino-2-phenylindole, DAPI (Sigma, UK) in 1x PBS for 10 min. Cells were kept in 1x PBS at 4°C until the cell images were visualized on an inverted fluorescence Axio Observer D1 microscope (Zeiss, UK). Four random image fields from each cell culture well were captured at a magnification of x100 by an AxioCam MR3 combined with ZEN Imaging software (Zeiss, UK). The cell fusion index was then evaluated as the number of nuclei in MF20-positive myotubes containing ≥ 3 nuclei and expressed as the percentage of the total nuclei number in the image field.

### Animals

Animals were bred in a minimal disease facility at Royal Holloway University of London, with food and water *ad libitum*. FLExDUX4 (JAX 028710) and HSA-MCM (JAX 025750) mice were purchased from The Jackson Laboratory (Maine, USA). FLExDUX4 colony was maintained as homozygous for Gt(ROSA)26Sor^tm1.1(DUX4*)Plj^ while HSA-MCM colony was maintained as hemizygous for Tg(ACTA1-cre/Esr1*)2Kesr. Tamoxifen (TMX)-inducible Cre-driver FLExDUX4 bi-transgenic line (aka. MCM-D4) used in this study was generated by crossing FLExDUX4 females with HSA-MCM males. Due to gender specific-DUX4 pathology in the MCM-D4 model, only males were used and littermates were allocated equally between groups. All mice were kept under a standard 12-hour light/dark cycle and were supervised on a daily basis by experienced animal staff.

### *In vivo* experimental design

In the initial study optimizing tamoxifen (TMX) dosage for inducing constant *DUX4* expression, 16-week-old MCM-D4 mice received either a single dose of 5 mg/kg TMX (*n*=5) or 2.5 mg/kg TMX once every 2 weeks (*n*=5) via intraperitoneal (IP) injection. TMX (Sigma, UK) was prepared as described previously (29) and diluted in warmed sterile corn oil to 1 mg/ml prior to use. HSA-MCM mice, considered as wild-type control (*n*=5), received volume-match corn oil. Bodyweight was recorded weekly. *In situ* TA force measurement was performed at week 4 while mice were under terminal anesthesia prior to TA muscle dissection.

In the subsequent study evaluating antisense therapeutic effect, 16-week-old MCM-D4 males were IP injected with 2.5 mg/kg TMX on days 0 and 14 to induce *DUX4* expression. HSA-MCM mice receiving the same TXM dosage were considered as wild-type control (*n*=4). MCM-D4 mice were further IP injected with 10 mg/kg of Vivo-PMO PACS4 (*n*=5), or 10 mg/kg of Vivo-PMO SCR (*n*=4), considered as a negative control, on days 2, 8, 16, 22 while HSA-MCM mice received volume-matched saline. Animals underwent treadmill exhaustion tests (days −6, 11, 25) and locomotor behavioral tests (days 22, 25, 28) prior to being put under terminal anesthesia for *in situ* TA force measurement and subsequent tissue collection (day 30). Mice were kept under isoflurane-induced anesthesia (3% in 100% CO_2_) during injections and were continuously monitored until they fully recovered, and then hourly for 3 hours post recovery.

### Treadmill exhaustion test

All tests were performed on a Treadmill Simplex II system (Columbus Instruments, Ohio, USA), with adjusted 15°C inclination. Mice were acclimatized to the apparatus for 5 min on an unmoving treadmill and at a speed of 5 m/min for additional 5 min. The speed was then increased by 0.5 m every minute. Mice were exercised until they were unable to remain off the stopper for 10 sec. Total running time was recorded and displayed as time to fatigue as percentage of the baseline time recorded on day −6.

### Open-field locomotor activity

Mouse open-field behavioral activity was examined using locomotor activity monitors as previously described (63). Mice were acclimatized to the test chamber during an undisturbed period of 15 min before the data were acquired and collected by Amon Lite software (version 1.4) every 10 min in a 60-min session. The same procedure was repeated every 3 days for 3 times. Data obtained from each mouse were averaged. During the acquisition, particular care was taken to minimize noise and movement into the room. Both the locomotor activity monitors and the software were obtained from Linton Instrumentation, UK.

### *In situ* muscle force measurement

Mice were anesthetized by IP injection with a mixture of 10 mg/ml dolethal (Vetoquinol, UK) and 15 μg/ml buprenodale (Dechra, UK) at 5 times of the bodyweight. The distal tendon of tibialis anterior (TA) muscle was dissected and attached to an isometric transducer, Dual-mode muscle lever (Aurora Scientific, Canada), through a loop made of braided silk suture (Harvard Apparatus, UK). The sciatic nerve was isolated and distally stimulated by a bipolar silver electrode using supramaximal square wave pulses at 0.1 ms duration. Data provided by the isometric transducer were recorded and analyzed using Dynamic Muscle Control and Analysis Software (Aurora Scientific, Canada). All isometric measurements were obtained at an initial length at which a maximal tension was recorded during the tetanus. Responses to tetanic stimulations at increased pulse frequencies from 10 Hz to 180 Hz were recorded and the maximal force (mN) was determined. The specific force (mN/mm^2^) was subsequently calculated based on a ratio of the maximal force and the muscle cross-sectional area (CSA) that was approximated mathematically by dividing the muscle mass by the optimum fiber length and the density of mammalian muscle, as described in TREAT-NMD SOP DMD_M.2.2.005. Force measurement was performed in a blinded manner.

### Post-mortem tissue processing

From each mouse, the gastrocnemius (GAS), quadriceps (QUAD) and TA muscles were collected. Tissues from one side of hindlimb were frozen immediately in liquid nitrogen for RNA extraction (as described above). Contralateral muscles were embedded in optimal cutting temperature medium (VWR, UK) and subsequently frozen in liquid nitrogen-cooled isopentane (Sigma, UK) and stored at −80°C. Frozen TA muscle was cryosectioned on an OTF5000 cryostat (Bright, UK) at 10-μm thickness for 10 serial levels through the muscle length, and transverse sections were collected onto SuperFrost slides.

### Immunohistochemistry staining

For collagen immunostaining, serial muscle section-containing slides were fixed in ice-cold acetone for 10 min and blocked in 1% (w/v) BSA, 1% (v/v) goat serum, 0.1% (v/v) Triton X-100 and 1x PBS for 1 hr. Subsequent incubation with rabbit anti-collagen VI (1:300, Abcam, UK) antibodies was carried out at 4°C, overnight. Slides were washed three times in 1x PBS, 0.05% (v/v) Tween-20 prior to 1-hr incubation with goat anti-rabbit AlexaFluor488 (1:1000, Life Biotechnologies, UK). An additional incubation for 15 min with 1 μg/ml DAPI was performed before slides were mounted in Mowiol 4-88. Reagents were purchased from Sigma, UK unless stated otherwise. Images from the largest mid-belly muscle sections were captured on Axio Observer D1 fluorescence microscope (Zeiss, UK) at a magnification of x100 by an AxioCam MR3 and were automatically stitched together by ZEN Imaging software (Zeiss, UK) to generate an image of the whole transverse muscle section.

For laminin and myosin heavy chain (MyHC) co-immunostaining, frozen sections were fixed in ice-cold acetone for 10 min, then blocked in mouse-on-mouse blocking buffer (Vector Laboratories, UK) supplemented with 1% (w/v) BSA, 1% (v/v) goat serum, 0.1% (v/v) Triton X-100, and 1x PBS for 30 min. Subsequent incubation with primary antibodies was carried out at 4°C for overnight, and then with compatible secondary antibodies for 1 hr, at room temperature. Primary antibodies were rabbit anti-laminin antibody (1:300, Abcam, UK) and mouse anti-MyHC antibodies (DSHB, Iowa, USA), including SC-71 and BF-F3 for fast twitch fibers type IIA and IIB, respectively (1:5), BF-G6 for embryonic MyHC (1:50), unstained fibers were considered as IIX. All DSHB antibodies were deposited by Schiaffino, S. Secondary antibodies (Life Technologies, UK) were goat anti-rabbit IgG Alexa568 (1:400) or goat anti-rabbit IgG Alexa488 (1:400), goat anti-mouse IgG2b Alexa488 (1:400), goat anti-mouse IgM Alexa405 (1:250), and goat anti-mouse IgG Alexa568 (1:400), respectively. Nuclei were stained with 1 μg/ml DAPI. Slides were mounted in Mowiol 4-88 (Sigma, UK). Images of whole transverse muscle sections were acquired and generated as described above.

### Histological analyses

Laminin staining was used for identifying the fiber perimeter. The number of total myofibers, MyHC-IIA, −IIB, or −IIX fibers, as well as the minimal Ferret’s diameter of individual fibers was automatically measured by MuscleJ software (National Institutes of Health, Maryland, USA) while the number of eMyHC was counted manually. Automatic analysis of the frequency distribution of the minimal Ferret’s diameter was carried out using GraphPad Prism6 software (California, USA). The CSA of the entire muscle sections or the area positive with collagen staining was semi-automatically scored by MuscleJ software. Fibrotic area was calculated as the percentage of the total area of the muscle cross-section. All analyses were performed by a single operator.

### Statistical analysis

Data were analyzed using GraphPad Prism6 software (California, USA) and are shown as the means ± S.E.M. Error bars represent the S.E.M; “*n*” refers to the number of biological replicates in cell culture work or the number of mice per group. Comparisons of statistical significance were assessed by one- or two-way ANOVA followed by Tukey’s *post-hoc* test. Significant levels were set at \**p* < 0.05, \*\**p* < 0.01, \*\*\**p* < 0.001, \*\*\*\**p* < 0.0001.

### Study approval

Animal procedures were performed in accordance with the UK Animals (Scientific Procedures) Act, 1986.

## Author Contributions

N.L-N., L.P. and G.D. conceived and designed the study. N.L-N. and A.M. performed muscle functional tests. N.L-N. performed other analyses. All authors contributed to result interpretation. N.L-N. wrote the manuscript with input from A.M., L.P. and G.D. All authors read and approved the final manuscript.

## Acknowledgements

We thank Dr Vincent Mouly, Institute of Myology, France for providing us the FSHD immortalized myoblast clones, A5 and A10. We also thank Drs Takako Jones and Peter Jones, University of Nevada, Nevada, USA for valuable advice on maintaining the mouse colonies and for a detailed protocol on using tamoxifen to induce *DUX4* expression in MCM-D4 mice. This study was supported by the Muscular Dystrophy UK (grant reference number: 16GRO-PG36-0083). The funder is not involved in study design, data interpretation, or writing of the manuscript.

## Conflict of interest statement

A patent named “Antisense oligonucleotides and uses thereof” has been filed by Royal Holloway University of London, UK and Institute of Myology, Paris, France. L.P. and G.D. are named inventors. The other authors have declared that no conflict of interest exists.

