## Supplemental Data for "Systemic antisense therapeutics inhibiting *DUX4* expression improves muscle function in an FSHD mouse model"

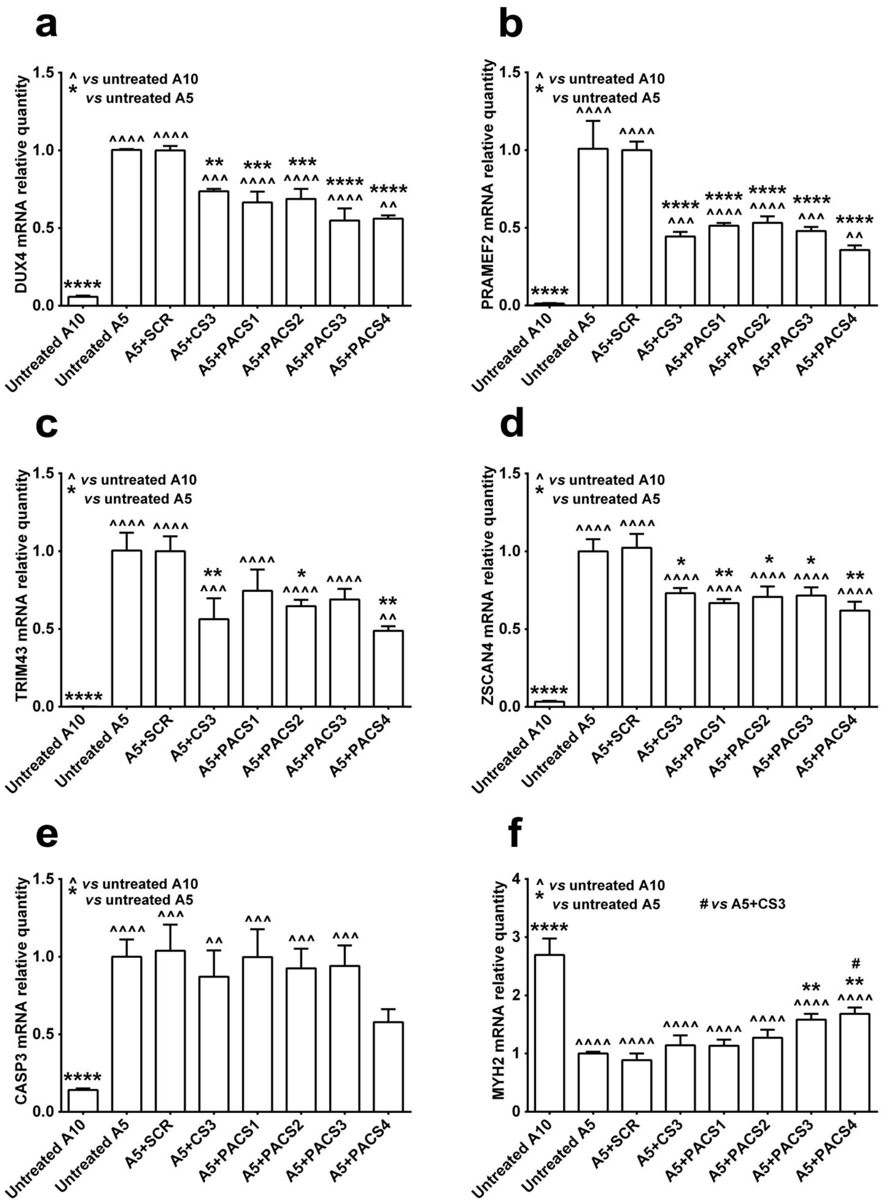


**Figure S1: Effect of 1 μM PMO treatment in FSHD immortalized myoblast cell cultures.** FSHD immortalized A5 myoblasts were differentiated for 2 days before the cells were treated with 1 μM PMOs via Endo-Porter-mediated transfection. Immortalized A10 or A5 cells receiving only Endo-Porter reagent were considered as untreated positive or negative control, respectively. Total RNA was extracted 2 days after PMO treatment. RT-qPCR quantification for *DUX4* **(a)** and its targets: *PRAMEF2* **(b)**, *TRIM43* **(c)**, *ZSCAN4* **(d)**, as well as markers of cell apoptosis *CASP3* **(e)** and cell differentiation *MYH2* **(f)** are shown. Statistical comparison was by one-way ANOVA followed by Tukey's multiple comparisons test, and was against untreated A10 or A5 cells, or A5 treated with PMO CS3 (considered as positive PMO control). Data are shown as mean ± S.E.M., $n = 3$, *$p < 0.05$, **$p < 0.01$, ***$p < 0.001$, ****$p < 0.0001$.

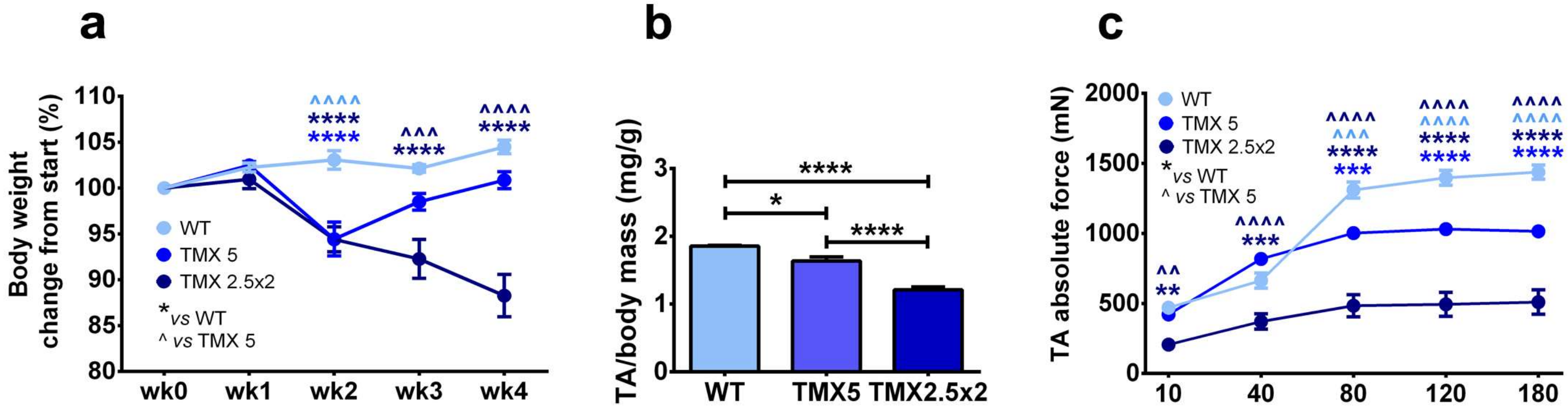


**Figure S2: Optimizing tamoxifen (TMX) dosage for inducing progressive DUX4 pathology in double transgenic MCM-D4 mice.** Male MCM-D4 mice received either a single dose of 5 mg/kg TMX (wk 0) or 2.5 mg/kg/biweekly TMX (wk0, wk2) via intraperitoneal (IP) injection. A group of HSA-MCM mice receiving volume-matched corn oil was considered as wild-type (WT) control. Body weight recorded weekly is displayed as the percentage of weight change from initial **(a)**. Tibialis anterior (TA) muscle mass normalized to the initial body weight is shown **(b)**. Four weeks after the first TMX injection, mice were put under terminal anesthesia for *in situ* TA force measurement; the absolute force is shown **(c)**. Statistical comparison was by one-way (b) or two-way (a, c, d) ANOVA followed by Tukey's multiple comparisons test. Data are shown as mean ± S.E.M., $n = 5$, *$p < 0.05$, **$p < 0.01$, ***$p < 0.001$, ****$p < 0.0001$.

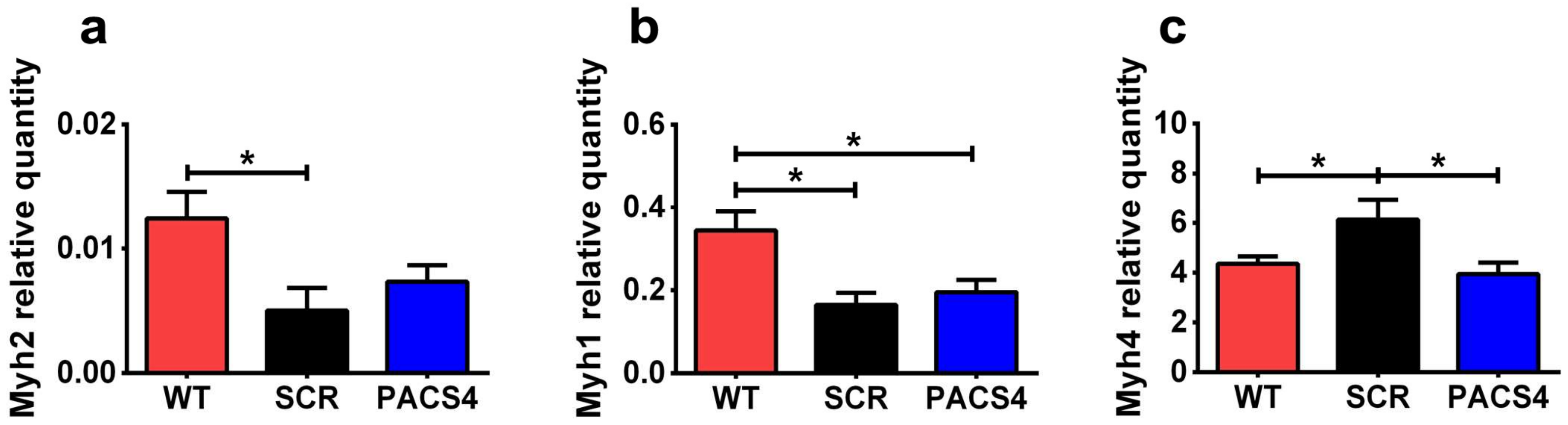


**Figure S3: Gene expression of major myofiber types in TA muscle.** mRNA levels of four fiber types were assessed by RT-qPCR, including *Myh2* for MyHC IIA **(a)**, *Myh1* for MyHC IIX **(b)**, and *Myh4* for MyHC IIB **(c);** *Myh7* for MyHC I was undetectable, relative to corresponding *Gapdh* expression. Data are shown as means ± S.E.M; $n$ = 4-5. Statistical comparison was by one-way ANOVA followed by Tukey's *post-hoc* test; $^*p < 0.05$.

**Table S1: Sequences of PMOs and targeting regions within *DUX4* 3'UTR**

| Name | PMO sequence (5'-3') | Target sequence (5'-3') |
|---|---|---|
| PMO SCR, 25-mer | CCTCTTACCTCAGTTACAATTTATA | |
| PMO CS3, 30-mer (+2 +31) | TATAGGATCCACAGGGAGG**A**GGCATTTTAA | TTAAAATGCC**C**CCTCCCTGTGGATCCTATA |
| PMO PACS1, 30-mer (-2 +28) | AGGATCCACAGGGAGGGGGCATTTTAATAT | ATATTAAAATGCCCCCTCCCTGTGGATCCT |
| PMO PACS2, 30-mer (-1 +29) | TAGGATCCACAGGGAGGGGGCATTTTAATA | TATTAAAATGCCCCCTCCCTGTGGATCCTA |
| PMO PACS3, 30-mer (-2 +28) | AGGATCCACAGGGAGG**A**GGCATTTTAATAT | ATATTAAAATGCC**C**CCTCCCTGTGGATCCT |
| PMO PACS4, 30-mer (-1 +29) | TAGGATCCACAGGGAGG**A**GGCATTTTAATA | TATTAAAATGCC**C**CCTCCCTGTGGATCCTA |
| PMO PACS4, 28-mer (+1 +28) | AGGATCCACAGGGAGG**A**GGCATTTTAAT | ATTAAAATGCC**C**CCTCCCTGTGGATCCT |

Polyadenylation signal (PAS): ATTAAA (target sequence) → TTTAAT (PMO sequence)

Cleavage site (CS): GATCCT (target sequence) → AGGATC (PMO sequence)

Modified nucleotide: **C** (target sequence) → **A** (PMO sequence)

**Table S2: Changes in mouse locomotor behavior after 4 weeks of Vivo-PMO treatment**

| Open-field cage activity | | WT | | SCR | | PACS4 | | SCR vs WT | PACS4 vs WT | PACS4 vs SCR |
|---|---|---|---|---|---|---|---|---|---|---|
| **Parameters** | **Descriptions** | **Mean** | **SEM** | **Mean** | **SEM** | **Mean** | **SEM** | ***p* value** | ***p* value** | ***p* value** |
| Total activity | Total beam breaks | 1633.2 | 132.6 | 267.5 | 55.0 | 702.2 | 79.2 | < 0.0001 | < 0.0001 | < 0.0001 |
| Fast activity | Fast beam breaks | 185.7 | 24.3 | 26.6 | 8.1 | 35.4 | 5.3 | 0.1638 | 0.2596 | 0.9747 |
| Slow activity | Slow beam breaks | 1447.5 | 118.8 | 240.9 | 47.6 | 666.8 | 74.7 | < 0.0001 | < 0.0001 | < 0.0001 |
| Total static counts | Total beam breaks (movement lower than mobile threshold) | 1233.0 | 93.4 | 222.6 | 40.0 | 629.2 | 71.6 | < 0.0001 | < 0.0001 | < 0.0001 |
| Fast static counts | Beam breaks (movement lower than mobile threshold and faster than fast threshold) | 62.5 | 9.4 | 8.1 | 1.6 | 18.9 | 3.3 | 0.9601 | 0.9656 | 0.9909 |
| Slow static count | Beam breaks (movement lower than mobile threshold and slower than fast threshold) | 1170.5 | 87.9 | 214.5 | 39.1 | 610.3 | 68.9 | < 0.0001 | < 0.0001 | < 0.0001 |
| Total mobile counts | Total beam breaks (movement greater than mobile threshold) | 400.2 | 47.1 | 44.9 | 19.6 | 73.0 | 12.1 | < 0.0001 | < 0.0001 | 0.9116 |
| Fast mobile counts | Beam breaks (movement greater than mobile threshold and faster than fast threshold) | 123.2 | 16.3 | 18.5 | 7.8 | 16.5 | 2.5 | 0.4654 | 0.4405 | 0.9985 |
| Slow mobile counts | Beam breaks (movement greater than mobile threshold and slower than fast threshold) | 277.0 | 36.2 | 26.4 | 11.9 | 56.5 | 9.7 | 0.0011 | 0.0026 | 0.9502 |
| Total rearing counts | Number of rearing beam breaks | 508.5 | 45.1 | 38.9 | 10.1 | 96.3 | 13.5 | < 0.0001 | < 0.0001 | 0.032 |
| Fast rearing counts | Number of fast rearing beam breaks | 241.6 | 26.1 | 25.3 | 7.3 | 47.0 | 7.3 | 0.0075 | 0.0107 | 0.9307 |
| Slow rearing counts | Number of slow rearing beam breaks | 266.9 | 27.0 | 14.4 | 3.1 | 49.3 | 6.9 | 0.0036 | 0.0279 | 0.9998 |
| Total centre rearing counts | Number of rearing beam breaks occurred away from the cage walls | 85.0 | 13.0 | 3.9 | 1.4 | 10.1 | 1.8 | 0.7508 | 0.8459 | 0.9985 |
| Fast centre rearing counts | Number of fast rearing beam breaks occurred away from the cage walls | 36.1 | 4.9 | 2.6 | 1.1 | 4.1 | 1.1 | 0.8722 | 0.9566 | > 0.9999 |
| Slow centre rearing counts | Number of slow rearing beam breaks occurred away from the cage walls | 48.2 | 9.7 | 1.0 | 0.4 | 6.0 | 0.9 | 0.8722 | 0.9566 | 0.9995 |

| | | | | | | | | | | |
|---|---|---|---|---|---|---|---|---|---|---|
| Active time | Time of mobile or static activity (sec) | 1153.5 | 86.4 | 215.4 | 41.3 | 584.5 | 64.0 | < 0.0001 | < 0.0001 | < 0.0001 |
| Static time | Time of static activity (sec) | 943.8 | 68.1 | 189.7 | 33.7 | 538.5 | 59.7 | < 0.0001 | < 0.0001 | < 0.0001 |
| Mobile time | Time of mobile activity (sec) | 209.7 | 24.1 | 25.8 | 10.3 | 45.9 | 7.1 | < 0.0001 | < 0.0001 | 0.9589 |
| Rearing time | Time spent rearing (sec) | 947.4 | 92.9 | 59.9 | 14.3 | 182.6 | 18.1 | < 0.0001 | < 0.0001 | 0.0161 |
| Front to back counts | Number of traverses from front to back | 115.2 | 11.4 | 18.3 | 5.2 | 35.5 | 5.7 | < 0.0001 | < 0.0001 | 0.0408 |
| Inactive time | Time spent in inactivity (sec) | 2446.5 | 86.4 | 3384.6 | 41.3 | 3015.5 | 64.0 | < 0.0001 | < 0.0001 | < 0.0001 |
| Distance travelled meters | Total distance travelled (m) | 72.8 | 6.0 | 14.6 | 3.1 | 31.8 | 3.6 | < 0.0001 | < 0.0001 | 0.0238 |

Data were assessed by GraphPad Prism6 (California, USA). Statistical significance was analyzed by one-way ANOVA followed by Tukey's *post-hoc* test.

**Table S3: Details of qPCR primers**

| Target gene | Accession number | Primer sequence (5'-3') | Amplicon size (bp) | Annealing temp (°C) |
|---|---|---|---|---|
| *B2M* | NM_004048 | Forward: CTCTCTTTCTGGCCTGGAGG<br>Reverse: TGCTGGATGACGTGAGTAAACC | 67 | 60 |
| *CASP3* | NM_001354784 | Forward: ACTGGACTGTGGCATTGAGA<br>Reverse: GCACAAAGCGACTGGATGAA | 162 | 60 |
| *DUX4 3'UTR* | Gene ID: 100288687 | Forward: CTCTGTGCCCTTGTTCTTC<br>Reverse: TCCAGGAGATGTAACTCTAATCCA | 98 | 60 |
| *MYH2* | NM_017534 | Forward: TCAGGTCTTCCCCATGAACC<br>Reverse: GCTTATACACAGGCAGCCAC | 185 | 60 |
| *PRAMEF2* | NM_023014 | Forward: ACCTTCTTCAGTGGGCACCT<br>Reverse: TGGGAACTGGGAGAGACACT | 120 | 60 |
| *TRIM43* | NM_138800 | Forward: ACCCATCACTGGACTGGTGT<br>Reverse: CACATCCTCAAAGAGCCTGA | 100 | 60 |
| *ZSCAN4* | NM_152677 | Forward: CTGGAGCAGTTTATGATTGG<br>Reverse: AGCTTCCTGTCCCTGCATGT | 162 | 60 |
| *Casp3* | NM_001284409 | Forward: GAGCAGCTTTGTGTGTGTGA<br>Reverse: GGCAGGCCTGAATGATGAAG | 158 | 60 |
| *Col1α1* | NM_007742 | Forward: GAAACTTTGCTTCCCAGATGTC<br>Reverse: AGACCACGAGGACCAGAA | 94 | 58 |
| *Frg1* | NM_013522 | Forward: TGCAATTGGGCCCAGAGAACAATG<br>Reverse: GGCCAGTAAAGCCATCTTCCCATC | 64 | 60 |
| *Gapdh* | NM_008084 | Forward: TCCATGACAACTTTGGCATTG<br>Reverse: TCACGCCACAGCTTTCCA | 103 | 60 |
| *Myh1* | NM_030679 | Forward: GTGGAAGCTATCAAGGGTCTGC<br>Reverse: TCTTGCGGTCTTCCTCAGTTTG | 79 | 60 |
| *Myh2* | NM_001039545 | Forward: ACCCTCTTATTTCCCAGCTGCAC<br>Reverse: ACTGCTGAACTCACAGACCCTTAC | 61 | 60 |
| *Myh3* | NM_001099635 | Forward: ACCTCTAGCCGGATGGT<br>Reverse: AATTGTCAGGAGCCACGAAAAT | 103 | 60 |
| *Myh4* | NM_010855 | Forward: AGAAACTGGAGGCTAGGGTGAG<br>Reverse: TCGTGCTTACGAAGACCCTTGAC | 94 | 60 |
| *Myh7* | NM_001361607 | Forward: ATACGCATGCTTGTGCCGTAGG<br>Reverse: TTCCTTTCTCGGAGCCACCTTG | 65 | 60 |
| *Pax7* | NM_011039 | Forward: CTCAGTGAGTTCGATTAGCCG<br>Reverse: AGACGGTTCCCTTTGTCGC | 144 | 60 |

| | | | | |
|---|---|---|---|---|
| *Tgfβ1* | NM_011577 | Forward: TGACGTCACTGGAGTTGTACGG<br>Reverse: TCGAAAGCCCTGTATTCCGTCTC | 61 | 62 |
| *Trim36* | NM_178872 | Forward: TGAAAGTGGGAGTTGCTTCC<br>Reverse: GAATCAAAACAGGCGTCCTC | 127 | 60 |
| *Wfdc3* | NM_027961 | Forward: CTTCCATGTCAGGAGCTGTG<br>Reverse: ACCAGGATTCTGGGACATTG | 134 | 58 |
